# Targeting Cholesterol-Dependent Piezo1 Activation Impairs Amoeboid Migration in Melanoma Cells

**DOI:** 10.1101/2025.07.11.664494

**Authors:** Sylvia Kuang, Alleah Abrenica, Neelakshi Kar, Jeremy S. Logue

## Abstract

Bleb-based migration enables cancer cells to navigate the heterogeneous tumor microenvironment. Here, we report a phenotypic screen identifying drugs that inhibit bleb formation, a driver of amoeboid migration. Statins, including Fluvastatin and Pitavastatin, suppress amoeboid migration of melanoma cells in confined environments by reducing intracellular cholesterol. This disrupts plasma membrane tension sensing by Piezo1, lowering intracellular Ca^2^+ levels. Both cholesterol supplementation and Piezo1 activation rescue migration in confined environments, confirming their functional link. Notably, high cholesterol biosynthesis enzyme levels correlate with reduced patient survival in melanoma. These findings reveal that cholesterol is essential for confinement sensing through Piezo1, identifying cholesterol biosynthesis or uptake as rational therapeutic targets against metastasis.

**Significance Statement:** This study builds on a phenotypic drug screen that identified statins as inhibitors of bleb-based migration, a key mode of cancer cell movement through confined spaces. We show that statins reduce membrane cholesterol, disrupting the function of the mechanosensitive channel Piezo1 and impairing melanoma cell migration. Restoring cholesterol or activating Piezo1 rescues this effect, revealing a functional link between cholesterol and confinement sensing. Our findings highlight cholesterol biosynthesis as essential for invasive cell behavior and identify it as a therapeutic vulnerability. Importantly, elevated cholesterol pathway activity correlates with reduced survival in melanoma patients, underscoring the clinical relevance of targeting this pathway to limit metastasis.

## Introduction

Cell migration is a fundamental biological process for numerous physiological and pathological functions. Fibroblasts typically display mesenchymal migration, which involves integrin-dependent adhesion and actin polymerization-based protrusion at the leading edge (1). However, some cells are known to switch between mesenchymal and amoeboid migration modes to adapt to the mechanochemical properties of tissues.

Amoeboid migration operates independently of integrins and relies on plasma membrane blebs to drive protrusion at the leading edge (2). In cells with a high level of actomyosin contractility, bleb formation is driven by intracellular pressure and cytoplasmic flow (3). Bleb retraction occurs after the polymerization of new cortical actin and the recruitment of myosin, which typically happens rapidly (4). In melanoma, primary tumor cells promote local invasion by adopting an amoeboid phenotype (5). Under these conditions, a slow type of amoeboid migration is driven by the mechanical worrying of stromal type I collagen by blebs (6). Alternatively, cells may utilize large and stable blebs for rapid migration through mechanochemically challenging environments.

Under conditions of low adhesion and confinement, cells can adopt a fast amoeboid migration mode (7, 8). This mode involves forming a leader bleb, which is a large and stable bleb (9). Requiring only friction for force transmission, the rapid cortical actomyosin flow in leader blebs provides the motive force for cell movement (10). In addition to myosin, flow is sustained by a high level of actin severing at leader bleb necks (11). When vertically confined to 3 µm, numerous transformed cell types will adopt fast amoeboid (leader bleb-based) migration (7, 9). Although a phenotypically similar migration mode has been observed at the margins of melanoma and breast cancer tumors using intravital imaging, cells are more likely to adopt this mode under conditions of high planar confinement, like the peri-vascular/lymphatic space (12).

While the molecular mechanisms driving this phenotypic transition are not yet fully understood, Piezo1 has been identified as a key driver. In confined environments, cells contain high levels of intracellular Ca^2+^, which depends on plasma membrane tension sensing by Piezo1 (13). Recently, we demonstrated that intracellular Ca^2+^ activates INF2 to promote actin cytoskeletal remodeling, de-adhesion, and amoeboid (*i*.*e*., bleb-based) migration in confined environments (13). Specific regulators of actomyosin contractility (*e*.*g*., ROCK2) have also been identified as necessary for this phenotypic transition (13).

Beyond promoting invasion, blebs have been shown to recruit curvature-sensitive proteins (*e*.*g*., septins), which form oncogenic signaling hubs that contribute to anoikis resistance in melanoma cells (14). Thus, identifying active drugs against blebs may have significant therapeutic value. Here, we report the results of a phenotypic screen of approved drugs. We demonstrate that Fluvastatin inhibits amoeboid migration in confined environments. The activity of Fluvastatin depends on inhibiting HMGCR, a crucial enzyme within the cholesterol biosynthetic pathway (15). Remarkably, lowering intracellular cholesterol levels in melanoma cells was found to de-couple the response of Piezo1 to increasing levels of confinement. Thus, confinement sensing by cells requires cholesterol, which maintains lipid order to facilitate the transmission of plasma membrane tension to Piezo1.

## Results

Using a highly metastatic melanoma sub-line derived from A375 cells, cultured under non-adherent conditions, and treated with a low dose of Latrunculin (0.1 µM) to destabilize the cortical actin cytoskeleton and induce constitutive blebbing, we screened nearly 200 approved drugs for activity against blebbing by light microscopy (Fig. 1A) (16). The screen identified roughly a dozen molecules with activity against blebs, including Simvastatin, Fluvastatin, Sunitinib malate, and Sorafenib (Fig 1B). Interestingly, we previously demonstrated that the off-target binding of Simvastatin to Vimentin inhibits fast amoeboid (leader bleb-based) migration (17). Sunitinib malate and sorafenib are multi-kinase inhibitors (*e*.*g*., RAF, VEGFR) and are anticipated to indirectly impact blebbing through the many pathways in which they have activity. Fluvastatin is not known to bind Vimentin; therefore, we decided to determine if Fluvastatin can inhibit amoeboid (*i*.*e*., bleb-based) migration (18). We vertically confined cells to 3 µm using a PDMS ceiling coated with bovine serum albumin (BSA; 1%) to induce leader bleb-based migration (LBBM) (Fig. 1C-E & Video S1) (19). Relative to vehicle, *i*.*e*., control, Fluvastatin (10 µM) treated cells were ∼75% less likely to adopt a leader mobile (LM) phenotype (Fig. 1C-E&Video S2). Additionally, leader mobile cells treated with Fluvastatin were ∼50% slower, while directionality was unchanged (Fig. 1F-G). As cells encounter 2D and 3D confinement within tissues, we also determined if Fluvastatin impacts cells migrating through microchannels (8 * 8 * 100 µm; HWL) (20). In channels coated with VCAM-1 (1 µg/µL) to simulate the microvasculature, motile cells adopt a mesenchymal, amoeboid, or hybrid phenotype. Vehicle, *i*.*e*., control, treated cells predominantly adopt a hybrid phenotype (Fig. 1H-J). However, cells treated with Fluvastatin predominantly adopt a mesenchymal phenotype (Fig. 1H-J). Under conditions in which cells cannot adopt a mesenchymal phenotype (*i*.*e*., poorly adherent), Fluvastatin treatment reduces the number of motile cells by ∼50% (Fig. 1K-L). Thus, Fluvastatin selectively inhibits amoeboid migration.

**Figure 1.**
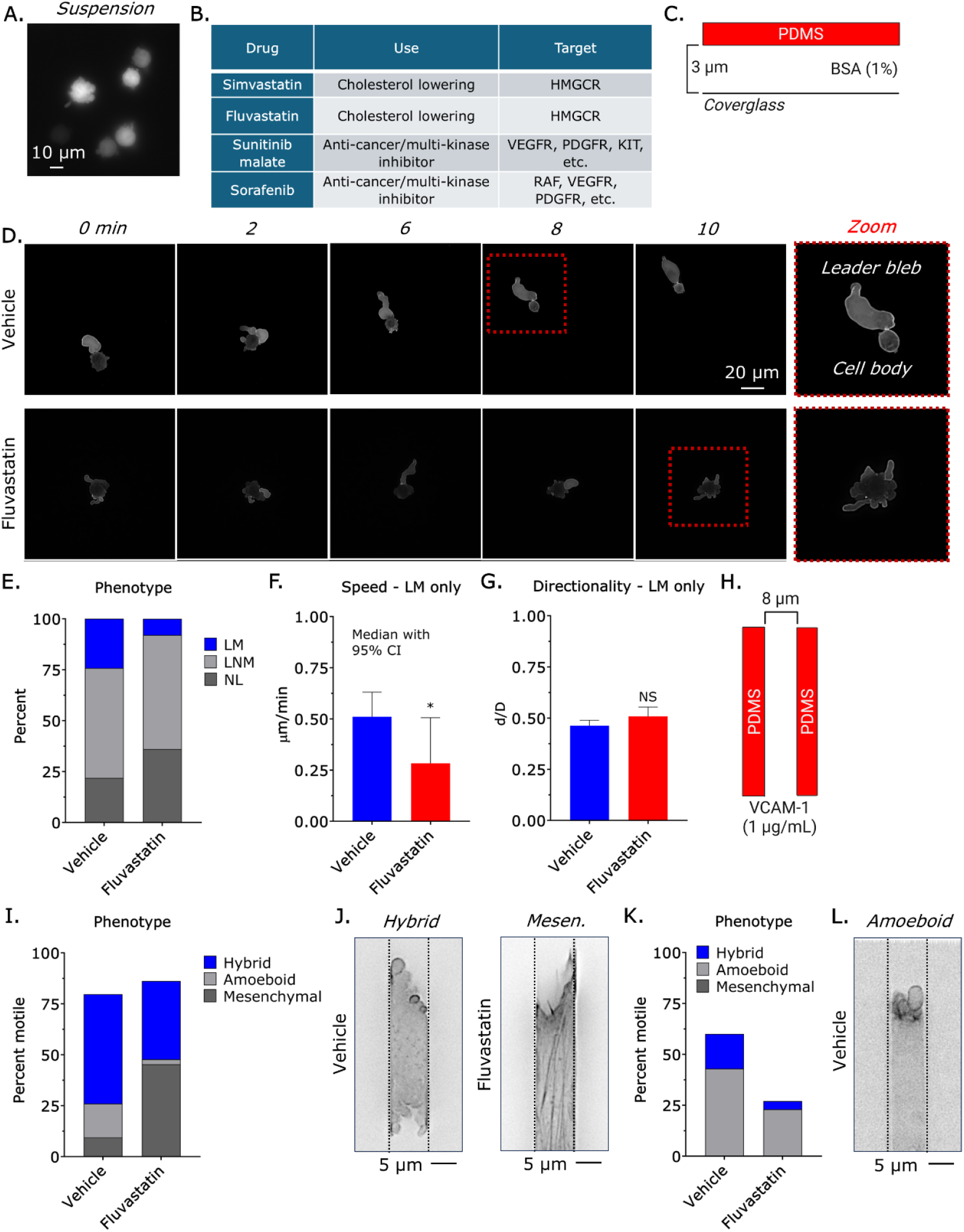
Fluvastatin selectively inhibits amoeboid migration. **A.** For phenotypic screening, we induced constitutive blebbing in A375 cells transduced with EGFP. **B**. Table of selected drugs identified in our phenotypic screen. **C**. Assay used to vertically confine cells down to ∼3 µm under non-adherent (BSA; 1%) conditions. **D**. Montage of vehicle (leader mobile; LM) and Fluvastatin (10 µM) treated (no leader; NL) cells. **E**. Percentage of leader mobile (LM), leader non-mobile (LNM), and no leader (NL) for vehicle (*n* = 297) and Fluvastatin (10 µM; *n* = 270) treated cells (χ^2^ < 0.0001). **F. - G**. Speed (median +/- 95% CI) and directionality (mean +/- SEM) for vehicle (*n* = 297) and Fluvastatin (10 µM; *n* = 270) treated leader mobile (LM) cells. **H**. Assay used for 3D confinement under adherent (VCAM-1; 1 µg/mL) conditions. **I**. Percentage of vehicle (*n* = 125) or Fluvastatin (10 µM; *n* = 170) treated cells adopting a hybrid, amoeboid, or mesenchymal phenotype in VCAM-1 (1 µg/mL) coated microchannels (χ^2^ < 0.0001). **J**. Representative images of cells treated with vehicle (hybrid) and Fluvastatin (10 µM; mesenchymal) in VCAM-1 (1 µg/mL) coated microchannels. **K**. Percentage of vehicle (*n* = 90) or Fluvastatin (10 µM; *n* = 153) treated cells adopting a hybrid, amoeboid, or mesenchymal phenotype in BSA (1%) coated microchannels (χ^2^ < 0.0001). **L**. Representative image of a cell adopting an amoeboid phenotype in a BSA (1%) coated microchannel. LifeAct-mEmerald was used to mark F-actin. Significance levels: * - p ≤ 0.05, ** - p ≤ 0.01, *** - p ≤ 0.001, and **** - p ≤ 0.0001. See also Video S1-2 and source data 1.

In dense stromal tissue, melanoma cells may adopt an amoeboid phenotype to migrate through confining pores (12). Therefore, we determined whether Fluvastatin selectively inhibits transmigration. On filters with 5 µm diameter pores, treatment with Fluvastatin inhibited transmigration by >50% (Fig. 2A). Whereas transmigration through larger pores was less affected (Fig. 2A). Within the cholesterol biosynthetic pathway, HMGCR catalyzes the formation of mevalonate (MVA) (21). Therefore, we determined if supplementation with MVA (100 µM) could rescue transmigration in Fluvastatin treated cells. Interestingly, MVA supplementation did not rescue transmigration but rescued cell proliferation (Fig. 2B-C). In cells treated with Fluvastatin, we found that MVA supplementation could not fully restore cholesterol levels (Fig. 2D). Suggesting that transmigration through confining pores may be particularly sensitive to intracellular cholesterol levels. Treatment with a hydrophilic statin, like Pravastatin, did not affect transmigration, proliferation, or intracellular cholesterol levels, as it cannot cross the plasma membrane (Fig. 2B-D) (22). Therefore, we determined if cholesterol supplementation could rescue transmigration through confining pores. While concentrations of 0.01 and 0.1 mM had little effect, supplementation with a 1 mM concentration of cholesterol was found to rescue transmigration (Fig. 2E). In the absence of Fluvastatin, the addition of cholesterol at this concentration was found to inhibit transmigration by ∼50% and dramatically inhibit cell proliferation (Fig. 2C-F). Other lipophilic statins, like Pitavastatin, were also found to inhibit transmigration (Fig. 2G). We also found that Fluvastatin inhibited the transmigration of Yale University mouse melanoma (YUMM) 3.3 cells, designed to model human melanoma mutations in a controlled mouse background (Fig. 2H) (23). In human foreskin fibroblasts, which are not known to adopt an amoeboid phenotype, Fluvastatin did not affect transmigration but dramatically inhibited cell proliferation (Fig. 2I-J) (7). These data demonstrate that transmigration through confining pores requires a precise intracellular cholesterol level.

**Figure 2.**
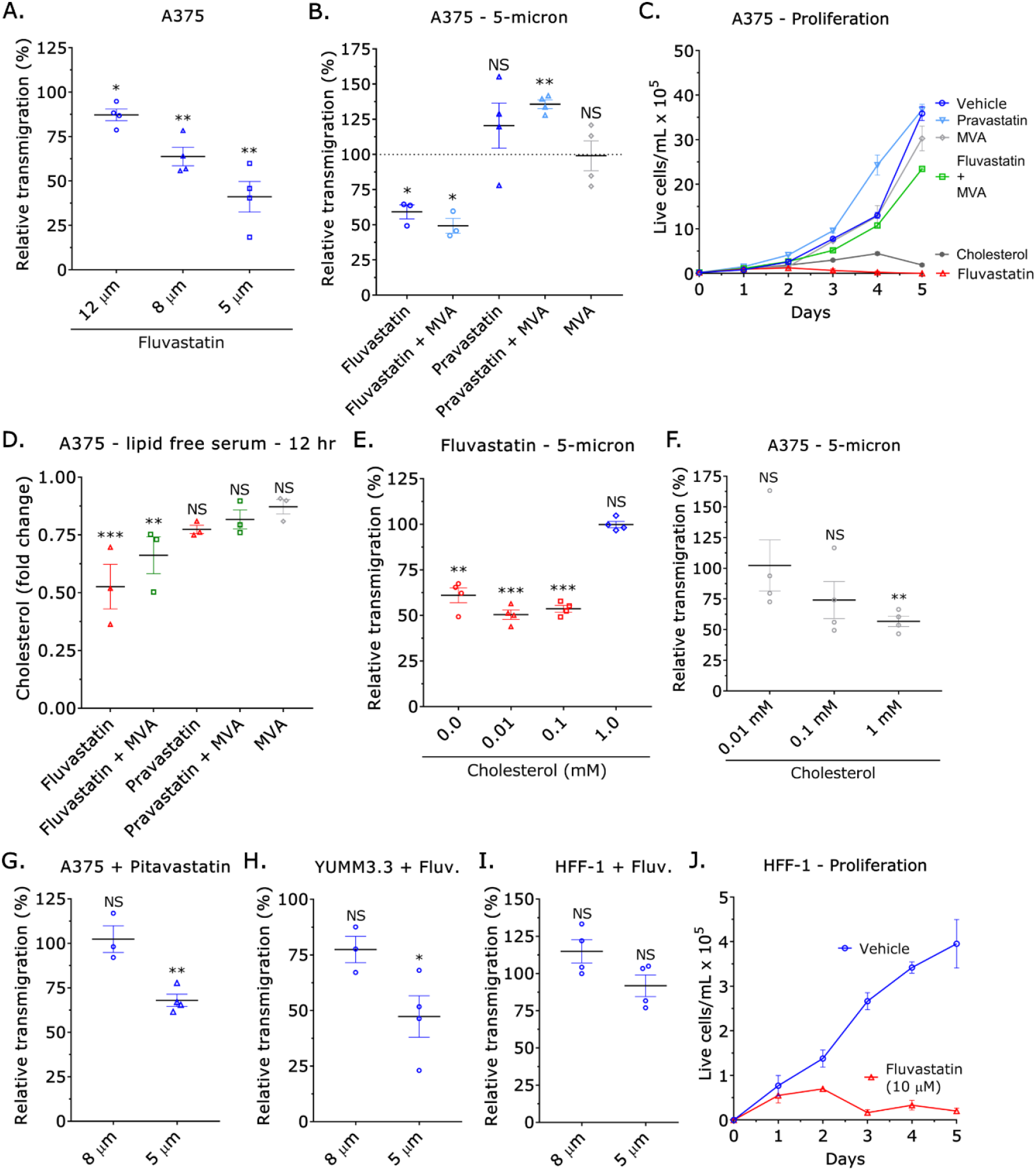
Intracellular cholesterol regulates transmigration through confining pores. **A.** Relative transmigration of Fluvastatin (10 µM) treated cells through 12, 8, or 5 µm pores towards fetal bovine serum (FBS; 10%) (mean +/- SEM). **B**. Relative transmigration of Fluvastatin (10 µM) +/- mevalonic acid (MVA; 100 µM), Pravastatin (10 µM) +/- mevalonic acid (MVA; 100 µM), and mevalonic acid (MVA; 100 µM) through 5 µm pores towards fetal bovine serum (FBS; 10%) (mean +/- SEM). **C**. Cell proliferation for vehicle, Fluvastatin (10 µM) +/- mevalonic acid (MVA; 100 µM), Pravastatin (10 µM) +/- mevalonic acid (MVA; 100 µM), mevalonic acid (MVA; 100 µM), and cholesterol (1 mM) over 5 days (mean +/- SEM). **D**. Fold change in cholesterol over 12 hr for Fluvastatin (10 µM) +/- mevalonic acid (MVA; 100 µM), Pravastatin (10 µM) +/- mevalonic acid (MVA; 100 µM), and mevalonic acid (MVA; 100 µM) treated cells cultured in the absence of serum lipids (mean +/- SEM). **E**. Relative transmigration of Fluvastatin (10 µM) treated cells supplemented with 0.01, 0.1, or 1.0 mM of cholesterol through 5 µm pores towards fetal bovine serum (FBS; 10%) (mean +/- SEM). **F**. Relative transmigration of cells supplemented with 0.01, 0.1, or 1.0 mM of cholesterol through 5 µm pores towards fetal bovine serum (FBS; 10%) (mean +/- SEM). **G**. Relative transmigration of Pitavastatin (10 µM) treated cells through 8 and 5 µm pores towards fetal bovine serum (FBS; 10%) (mean +/- SEM). **H**. Relative transmigration of Fluvastatin (10 µM) treated Yale University mouse melanoma (YUMM) 3.3 cells through 8 and 5 µm pores towards fetal bovine serum (FBS; 10%) (mean +/- SEM). **I**. Relative transmigration of Fluvastatin (10 µM) treated human foreskin fibroblast (HFF) 1 cells through 8 and 5 µm pores towards fetal bovine serum (FBS; 10%) (mean +/- SEM). **J**. Proliferation of vehicle and Fluvastatin (10 µM) treated human foreskin fibroblast (HFF) 1 cells over 5 days (mean +/- SEM). Significance levels: * - p ≤ 0.05, ** - p ≤ 0.01, *** - p ≤ 0.001, and **** - p ≤ 0.0001.

Statins, like Fluvastatin and Pitavastatin, inhibit HMGCR, the rate-limiting enzyme within the cholesterol biosynthetic pathway (24). Therefore, we determined if RNAi of HMGCR could similarly inhibit transmigration through confining pores. Relative to non-targeting, *i*.*e*., control, treatment with a siRNA towards HMGCR did not affect transmigration (Fig. 3A-B). However, intracellular cholesterol levels were unchanged even 3 days after treatment with an HMGCR siRNA (Fig. 3C). However, culturing cells in media containing lipid free serum led to significant reductions in intracellular cholesterol (Fig. 3D). Utilizing lipid free serum in transmigration assays, we observed a substantial decrease in transmigration (Fig. 3E). In agreement with these results, RNAi of HMGCR induced a compensatory increase in LDL uptake (Fig. 3F) (25). Although we observed a significant increase, Fluvastatin treatment did not upregulate LDL uptake to the same degree (Fig. 3G). Thus, the compensatory upregulation of LDL uptake induced by Fluvastatin treatment is insufficient to restore transmigration through confining pores. In agreement with these data, supplementation with cholesterol (1 mM) could rescue leader bleb-based migration (LBBM) in Fluvastatin treated cells (Fig. 3H).

**Figure 3.**
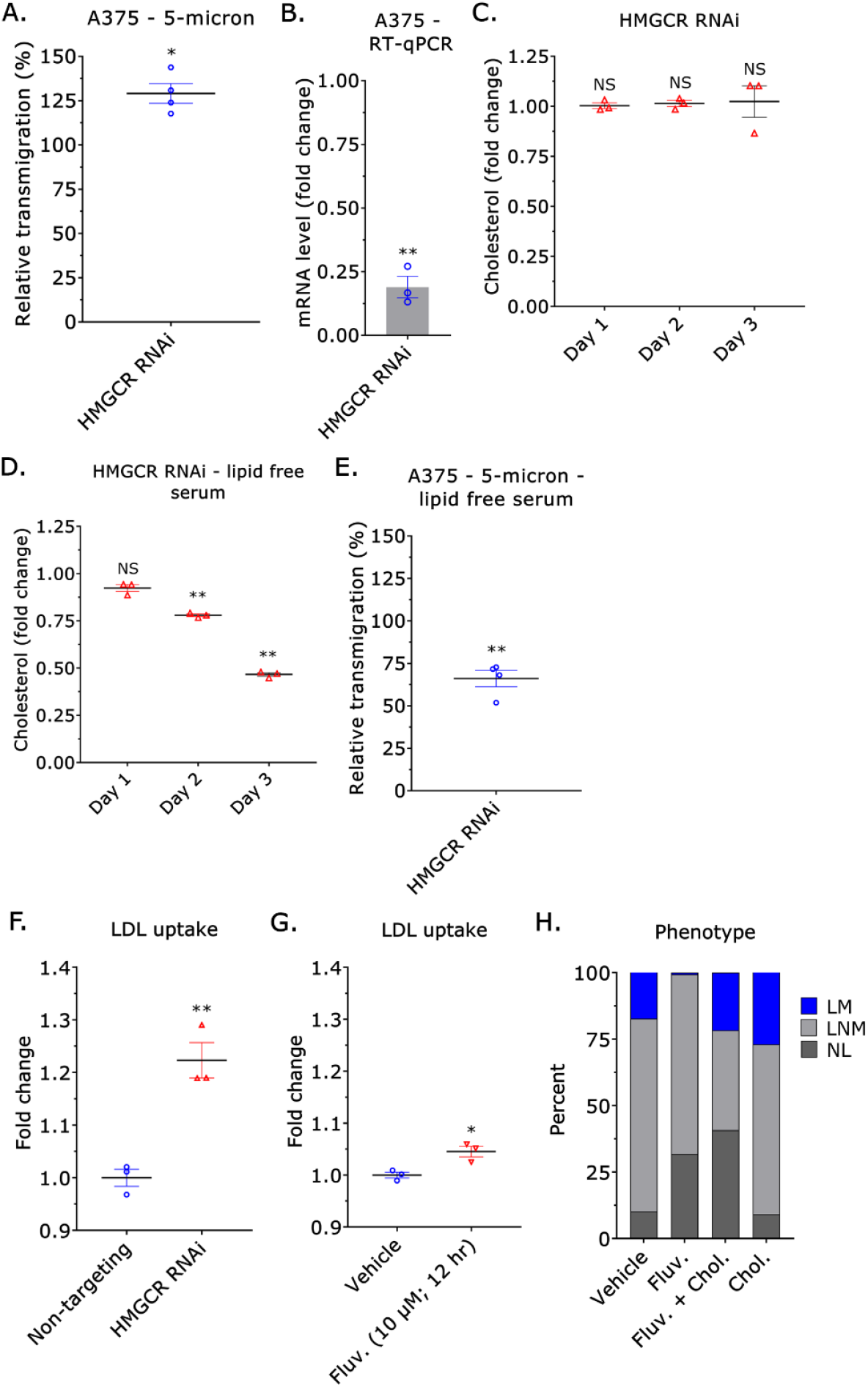
Fluvastatin modestly up-regulates LDL uptake. **A.** Relative transmigration of cells treated with an HMGCR siRNA through 5 µm pores towards fetal bovine serum (FBS; 10%) (mean +/- SEM). **B**. Fold change in mRNA after HMGCR RNAi, as measured by RT-qPCR (mean +/- SEM). **C**. Fold change in cholesterol after RNAi of HMGCR on days 1, 2, and 3 for cells cultured in the presence of serum lipids. **D**. Fold change in cholesterol after RNAi of HMGCR on days 1, 2, and 3 for cells cultured in the absence of serum lipids. **E**. Relative transmigration of cells treated with an HMGCR siRNA through 5 µm pores towards lipid-free serum (10%) (mean +/- SEM). **F**. Fold change in LDL uptake for cells treated with a non- targeting or HMGCR siRNA for 3 days (mean +/- SEM). **G**. Fold change in LDL uptake for cells treated with vehicle or Fluvastatin (10 µM) for 12 hr (mean +/- SEM). **H**. Percentage of leader mobile (LM), leader non-mobile (LNM), and no leader (NL) for vehicle (*n* = 156), Fluvastatin (10 µM; *n* = 161) (χ^2^ < 0.0001), Fluvastatin (10 µM) + cholesterol (1 mM; *n* = 47) (χ^2^ < 0.0001), and cholesterol (1 mM; *n* = 144) (χ^2^ = 0.03938) treated A375 cells transduced with EGFP. Significance levels: * - p ≤ 0.05, ** - p ≤ 0.01, *** - p ≤ 0.001, and **** - p ≤ 0.0001. See also source data 1.

Cholesterol is an essential component of cell membranes, modulates signaling (e.*g*., organizing lipid rafts), and serves as a precursor to biomolecules (*e*.*g*., steroid hormones) (26). As amoeboid (*i*.*e*., bleb-based) migration requires a high level of actomyosin contractility, we determined if upregulating actomyosin contractility could rescue leader bleb-based migration (LBBM). To accomplish this, we used RNAi to deplete cells of myosin phosphatase target subunit 1 (MYPT1), which is an essential component of myosin light chain phosphatase (MLCP) (7). In cells treated with Fluvastatin, we found that RNAi of MYPT1 had little effect on the number of leader mobile cells (Fig. S1A-B). By immunoblot, we also could not detect any changes in the level of phosphorylated regulatory light chain (pRLC; S19) (Fig. S1C). Additionally, we found that the response of ROCK1/2 to serum was not significantly different from cells pre-treated with vehicle (2 hr), as measured by a fluorescent biosensor (Fig. S1D) (27). In melanoma cells, these data demonstrate that low intracellular cholesterol levels do not interfere with actomyosin contractility.

We recently found that cell confinement increases intracellular [Ca^2+^] by activating the plasma membrane tension sensitive channel, Piezo1 (13). As cholesterol helps to maintain lipid order, we determined if supplementation could increase intracellular [Ca^2+^]. Using a genetically encoded Ca^2+^ indicator (GCaMP) to monitor transients (*i*.*e*., spontaneous oscillations in intracellular Ca^2+^ levels), we found that supplementation with 0.1 or 1 mM of cholesterol could increase the peak fold change by >3 (Fig. 4A-B) (28). We found that this effect depends on Piezo1, as an inhibitor or siRNA towards Piezo1 rendered cells unresponsive to cholesterol supplementation (Fig. 4C-E) (29). Fluvastatin treatment (10 µM; 12 hr) did not reduce the level of Piezo1 mRNA (Fig. 4F). Relative to non-targeting, *i*.*e*., control, treatment with a siRNA towards Piezo1 led to a significant decrease in transmigration through confining pores (Fig. 4G). Consistent with these findings, we found that intracellular [Ca^2+^] no longer correlated with confinement in cells treated with Fluvastatin, which could be rescued with cholesterol supplementation (Fig. 4H-K). Thus, cholesterol is found to be necessary for confinement sensing through Piezo1.

**Figure 4.**
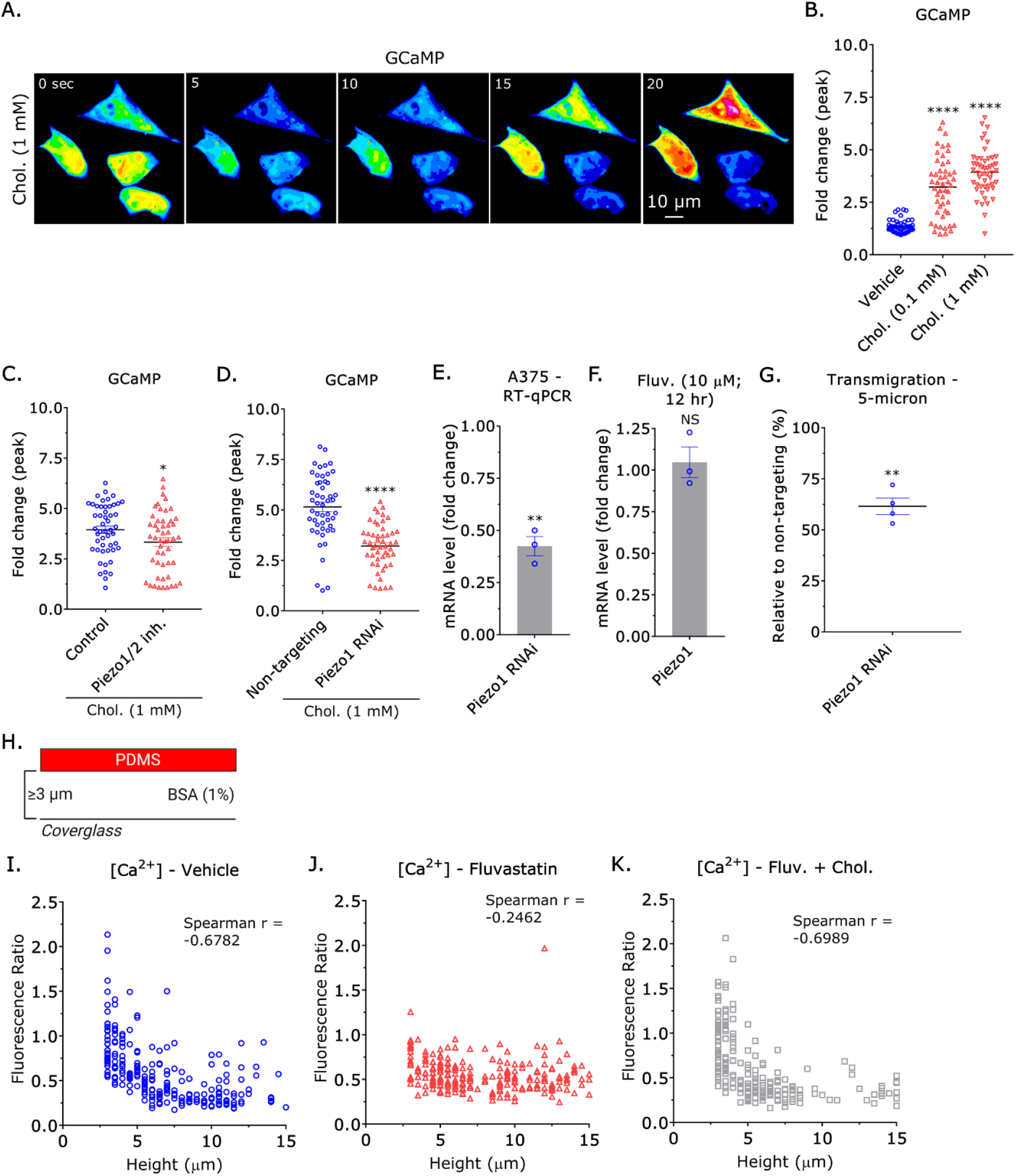
Cholesterol enables confinement sensing through Piezo1. **A.** Pseudocolored montage of cholesterol (1 mM) supplemented cells transfected with GCaMP. **B**. Quantification of spontaneous Ca^2+^ transients in cells treated with vehicle, 0.1, or 1 mM of cholesterol, as measured by the fold change in GCaMP fluorescence (mean +/- SEM). A Dunnett’s multiple comparison test was used to determine statistical significance. **C**. Quantification of spontaneous Ca^2+^ transients in cholesterol (1 mM) supplemented cells treated with a Piezo1/2 inhibitor (10 µM), as measured by the fold change in GCaMP fluorescence (mean +/- SEM). **D**. Quantification of spontaneous Ca^2+^ transients in cholesterol (1 mM) supplemented cells treated with a non-targeting or Piezo1 siRNA, as measured by the fold change in GCaMP fluorescence (mean +/- SEM). **E**. Fold change in mRNA after Piezo1 RNAi, as measured by RT-qPCR (mean +/- SEM). **F**. Fold change in Piezo1 mRNA after Fluvastatin treatment (10 µM; 12 hr), as measured by RT-qPCR (mean +/- SEM). **G**. Relative transmigration of cells treated with a Piezo1 siRNA through 5 µm pores towards fetal bovine serum (FBS; 10%) (mean +/- SEM). **H**. Assay used to variably (≥ 3 µm) confine cells under non-adherent (BSA; 1%) conditions. **I. - K**. Intracellular Ca^2+^ levels in vehicle, Fluvastatin (10 µM), and Fluvastatin (10 µM) + cholesterol (1 mM) treated variably confined (≥ 3 µm) cells under non-adherent conditions (BSA; 1%). We measured intracellular Ca^2+^ levels by taking the ratio of a red fluorescent Ca^2+^ indicator to a green fluorescent dye. Spearman rank correlation coefficients were used to determine statistical significance. Significance levels: * - p ≤ 0.05, ** - p ≤ 0.01, *** - p ≤ 0.001, and **** - p ≤ 0.0001. See also Figure S1.

Given that confinement sensing through Piezo1 requires cholesterol, we determined if an activator of Piezo1 could rescue migration in 2D and 3D environments. Fluvastatin treatment reduces the number of leader mobile cells by >75% (Fig. 5A-B). Additionally, leader mobile cells treated with Fluvastatin are ∼50% slower, while directionality is unchanged (Fig. 5C-D). However, we found that a Piezo1 activator (1 µM) could fully rescue fast amoeboid (leader bleb-based) migration (Fig. 5A-D) (30). In a 3D confined environment (*i*.*e*., microchannels), motile cells adopt either a mesenchymal, amoeboid, or hybrid phenotype. Vehicle, *i*.*e*., control, treated cells predominantly adopt a hybrid phenotype (Fig. 5E-F). However, cells treated with Fluvastatin predominantly adopt a mesenchymal phenotype (Fig. 5E-F). Adding a Piezo1 activator could rescue the hybrid phenotype (Fig. 5E-F). Additionally, we found that the Piezo1 activator triggered a ∼2-fold increase in the number of cells adopting an amoeboid phenotype (Fig. 5E-F). By maintaining lipid order, we propose that cholesterol promotes the transmission of plasma membrane tension to Piezo1, which increases intracellular [Ca^2+^], inducing a phenotypic switch to amoeboid (*i*.*e*., bleb-based) migration (Fig. 5G). Given that being able to switch between mesenchymal, amoeboid, and hybrid migration modes is anticipated to promote disease progression through multiple mechanisms, we correlated cholesterol biosynthesis enzymes with patient survival using transcriptomic data from The Cancer Genome Atlas (TCGA) for skin cutaneous melanoma (SKCM) (31). While we did not observe any significant changes in the level of any of these enzymes between primary and metastatic tumors, we did find that high levels of several enzymes, including mevalonate kinase (MVK), phosphomevalonate kinase (PMVK), isopentenyl-diphosphate delta isomerase 2 (IDI2), farnesyl diphosphate synthase (FDPS), and farnesyl-diphosphate farnesyltransferase 1 (FDFT1), were correlated with decreased rates of survival (p < 0.05) (Fig. 5H). Interestingly, higher levels of geranylgeranyl pyrophosphate synthase 1 (GGPS1) are associated with improved rates of survival, supporting the notion that the sterol and non-sterol branches of the mevalonate pathway compete for shared intermediates (*e*.*g*., farnesyl pyrophosphate) (Fig. 5H) (32). Therefore, high enzymatic activity within the sterol branch of the mevalonate pathway predicts a poor prognosis in melanoma.

**Figure 5.**
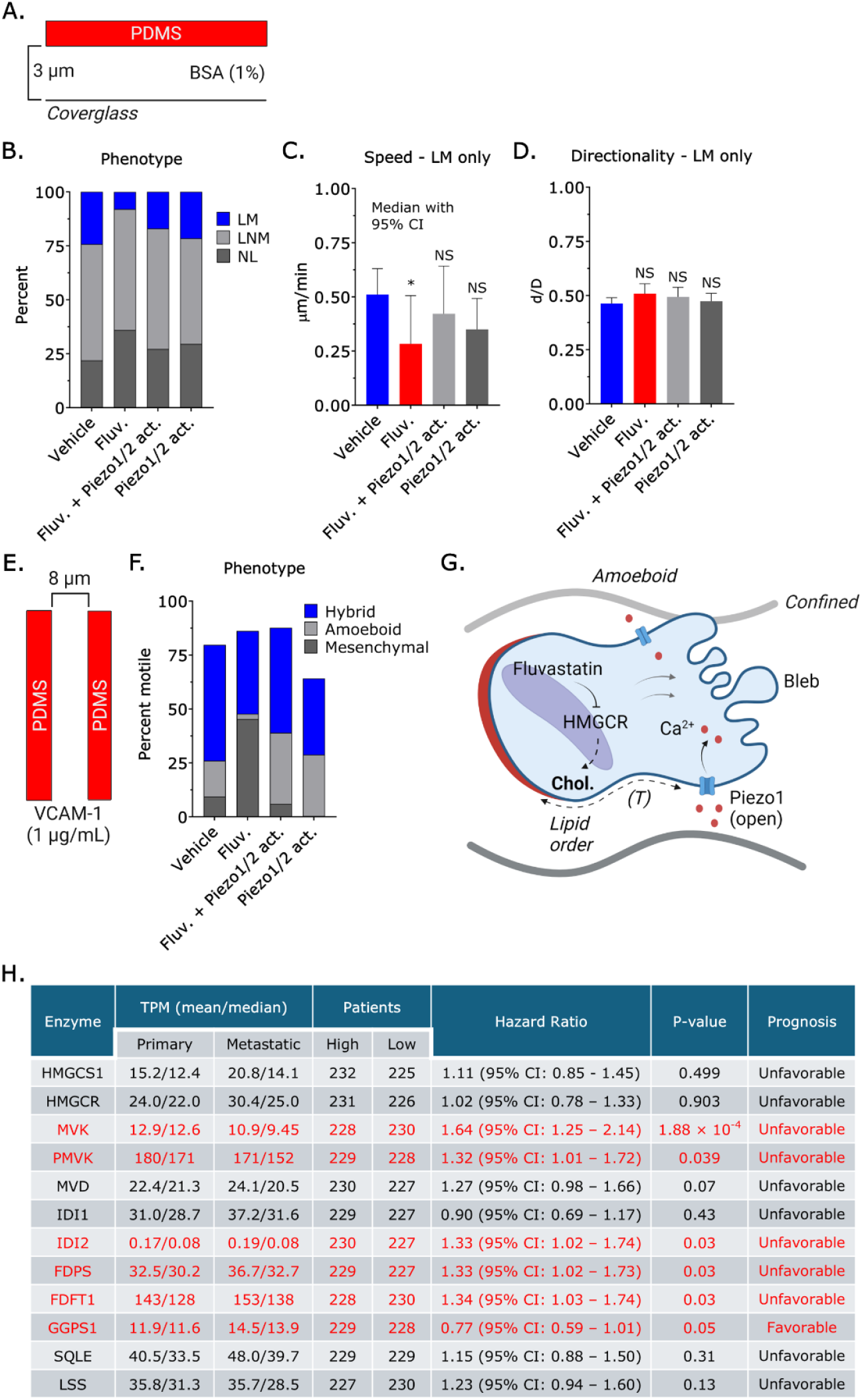
Piezo1 activation rescues migration defects caused by Fluvastatin. **A.** Assay used to vertically confine cells down to ∼3 µm under non-adherent (BSA; 1%) conditions. **B**. Percentage of leader mobile (LM), leader non-mobile (LNM), and no leader (NL) for vehicle (*n* = 297), Fluvastatin (10 µM; *n* = 270) (χ^2^ < 0.0001), Fluvastatin (10 µM) + Piezo1/2 activator (1 µM; *n* = 246) (χ^2^ = 0.4346), and Piezo1/2 activator (1 µM; *n* = 225) (χ^2^ = 0.1825) treated cells. **C. - D**. Speed (median +/- 95% CI) and directionality (mean +/- SEM) for vehicle (*n* = 297), Fluvastatin (10 µM; *n* = 270), Fluvastatin (10 µM) + Piezo1/2 activator (1 µM; *n* = 246), and Piezo1/2 activator (1 µM; *n* = 225) treated leader mobile (LM) cells. **E**. Assay used for 3D confinement under adherent (VCAM-1; 1 µg/mL) conditions. **F**. Percentage of vehicle (*n* = 125), Fluvastatin (10 µM; *n* = 170) (χ^2^ < 0.0001), Fluvastatin (10 µM) + Piezo1/2 activator (1 µM; *n* = 147) (χ^2^ = 0.0052), and Piezo1/2 activator (1 µM; *n* = 101) (χ^2^ < 0.0001) treated cells adopting a hybrid, amoeboid, or mesenchymal phenotype in VCAM-1 (1 µg/mL) coated microchannels. **G**. Model depicting the role of plasma membrane cholesterol in transmitting tension to Piezo1, promoting the transition to amoeboid (*i*.*e*., bleb-based) migration in confined environments. **H**. Table of cholesterol biosynthesis enzyme levels (transcripts per million; TPM) for primary and metastatic tumors, hazard ratios, and prognosis for skin cutaneous melanoma (SKCM). Significance levels: * - p ≤ 0.05, ** - p ≤ 0.01, *** - p ≤ 0.001, and **** - p ≤ 0.0001. See also source data 1.

## Discussion

The capacity to switch between mesenchymal, amoeboid, or hybrid migration modes is likely to promote disease progression through multiple mechanisms (33). In melanoma, the invasive front of primary tumors is characterized by rounded (*i*.*e*., amoeboid) cells (5). By intravital imaging, melanoma and breast cancer cells have been observed migrating in the direction of bleb formation (12). Src family kinase inhibitors, like Dasatinib, have also been shown to promote a switch from mesenchymal to amoeboid (*i*.*e*., bleb-based) migration (34). However, the switch to an amoeboid phenotype is most frequently observed under low adhesion and confinement (7). Accordingly, this phenomenon may help cancer cells migrate through the heterogeneous tumor microenvironment (20). Beyond promoting migration, bleb formation has also been implicated in resistance to anoikis (14). Therefore, phenotypic plasticity presents a significant barrier to therapeutic development.

Drug re-purposing offers several advantages, including faster development timelines, lower development costs, and improved safety profiles. Using constitutively blebbing melanoma cells, we found that Fluvastatin has activity against blebs. Although we identified additional drugs, like Sorafenib, we chose to focus on the statins, as they have a well established safety profile. Retrospective studies have also indicated that statin use increases survival rates (35). Using cells confined to 3 µm, which induces a phenotypic transition to fast amoeboid (leader bleb-based) migration, we found that Fluvastatin decreased the number of mobile cells (7-9). The assay simulates planar (*i*.*e*., 2D) confinement, including interstitial gaps, perivascular space, and between the basal surface of epithelial cells and the stroma (36, 37). Melanoma cells adopt a mesenchymal, amoeboid, or hybrid phenotype in other environments, like microvasculature (38). While vehicle, *i*.*e*., control, treated cells predominantly adopt a hybrid phenotype, Fluvastatin treated cells were predominantly mesenchymal in microfabricated channels coated with VCAM-1. In transmigration assays, cells migrating through the most confining pore sizes were most affected by Fluvastatin, which again highlights the specificity of Fluvastatin towards amoeboid (*i*.*e*., blebbing) cells. Human foreskin fibroblasts, which are not known to adopt an amoeboid phenotype, are also less affected by Fluvastatin. However, the proliferation of cancer and normal cells were both inhibited by Fluvastatin. We attribute this effect to a decrease in the prenylation of small GTPases; the mevalonate pathway produces intermediates important for prenylation (32). Small GTPases like Ras are essential for transmitting mitogenic (*i*.*e*., proliferative) signals. Additionally, cell migration may not require as much new membrane synthesis as proliferation.

Interestingly, while adding mevalonate (MVA) to Fluvastatin treated cells rescued proliferation, cell migration remained inhibited. We measured intracellular cholesterol levels and found that mevalonate supplementation did not fully restore them, suggesting that it may not be utilized as effectively as endogenously produced mevalonate. Adding cholesterol fully rescued migration, implying that amoeboid (*i*.*e*., bleb-based) migration is less dependent on protein prenylation. Without Fluvastatin, we found that additional cholesterol impairs cell migration and proliferation, likely by disrupting lipid raft-associated signaling (*e*.*g*., PI3K/Akt) (26). These data demonstrate that amoeboid migration requires a precise intracellular cholesterol level.

Cholesterol plays several critical roles in cells, including forming lipid rafts, serving as a precursor to steroid hormones, and maintaining lipid order (26). Whereas actomyosin contractility was unaffected, we found that cholesterol elevates [Ca^2+^] and that this effect depends on Piezo1, suggesting that its role in maintaining lipid order is essential for transmitting tension to Piezo1. Cholesterol maintains lipid order by filling voids between phospholipids, restricting fluidity (39). Recently, we demonstrated that Piezo1/Ca^2+^ activates INF2 to promote actin cytoskeletal remodeling, de-adhesion, and amoeboid (*i*.*e*., bleb-based) migration in confined environments (13). Here, Fluvastatin disrupted the Ca^2+^ response to confinement, which we could restore with cholesterol supplementation. Furthermore, a Piezo1 activator rescued amoeboid migration in cells treated with Fluvastatin. Thus, our data supports a model in which intracellular cholesterol enables cells to sense confined environments.

Although statins have pleiotropic effects, our work demonstrates that lipophilic statins, like Fluvastatin, are likely to have direct therapeutic effects on cancer cells by impairing intracellular cholesterol metabolism. Unlike RNAi of HMGCR, which led to compensatory LDL uptake, Fluvastatin treated cells displayed limited uptake, suggesting an insufficient compensatory response that may stem from the shorter duration of treatment. Simvastatin induces Vimentin bundling, raising the possibility that Fluvastatin may impair LDL uptake through cytoskeletal disruption (17, 18). Ultimately, the capacity of cells to upregulate LDL uptake may predict their sensitivity to statins. Collectively, our work identifies cholesterol biosynthesis or uptake as rational therapeutic targets against amoeboid migration, a key driver of melanoma progression.

## Resource Availability

### Lead contact

Requests for further information and resources should be directed to and will be fulfilled by the lead contact, Jeremy S. Logue.

### Materials availability

All unique/stable reagents generated in this study are available from the lead contact upon request.

### Data and code availability

- All data reported in this paper will be shared by the lead contact upon request.
- This paper does not report original code.
- Any additional information required to reanalyze the data reported in this paper is available from the lead contact upon request.

## Acknowledgments

We thank the Cady Lab (SUNY Polytechnic Institute, Albany, NY) for fabricating silicon wafer molds. This work was supported by grants from the Melanoma Research Alliance (MRA; award no. 688232) (DOI: https://doi.org/10.48050/pc.gr.91570), the American Cancer Society (ACS; award no. RSG-20-019-01 - CCG), and the National Institutes of Health (NIH; award no. R35GM146588) to J.S.L.

## Author Contributions

S.K.: Investigation, A.A.: Investigation, N.K.: Investigation, J.S.L.: Conceptualization, Supervision, Project administration, Visualization, Writing - Original Draft, Funding acquisition

## MATERIALS AND METHODS

### Cell lines

A375-M2 (CRL-3223), YUMM3.3 (CRL-3365), and HFF-1 (SCRC-1041) were obtained from the American Type Culture Collection (Manassas, VA). Cells were cultured in high-glucose DMEM supplemented with fetal bovine serum (FBS; 10%) (Sigma Aldrich; 12106C), L-glutamine (Thermo Fisher; 35050061), pyruvate, HEPES, and Antibiotic-Antimycotic (Thermo Fisher; 15240096).

### Drug Screen

A library of Federal Drug Administration (FDA) approved drugs (7200) was purchased from Tocris Bioscience (Bristol, UK). For screening, A375 cells transduced with EGFP (Addgene; 19070) were cultured in BSA (1%) coated glass-bottom plates (Cellvis; P96-1.5H-N). In addition to the drug, cells were treated with Latrunculin (0.1 µM) (Tocris Bioscience; 3973) for 1 hr before imaging. Analyze Particles in Fiji (https://fiji.sc/) was used to obtain shape descriptors.

### Plasmids

LifeAct-mEmerald (plasmid no. 54148) and GCaMP (plasmid no. 40753) were obtained from Addgene (Watertown, MA). Eevee-ROCK was kindly provided by Dr. Michiyuki Matsuda (Kyoto University). 1 µg of plasmid was used to transfect 750,000 cells in each well of a 6-well plate using Lipofectamine 2000 (5 µL; Thermo Fisher) in OptiMEM (400 µL; Thermo Fisher). After 20 min at room temperature, plasmid in Lipofectamine 2000/OptiMEM was incubated with cells in complete media (2 mL) overnight.

### Chemical treatments

Latrunculin (3973), Fluvastatin (3309), Pitavastatin (4942), Pravastatin (2318), Piezo1/2 inhibitor (4912), and Piezo1/2 activator (5586) were purchased from Tocris Bioscience (Bristol, UK). Mevalonic acid lactone (M4667) and water soluble cholesterol (C4951) were purchased from Sigma Aldrich (St. Louis, MO). Before confinement, cells were treated with drug for 1 hr. In parallel, confining devices were incubated with drug in complete media for at least 1 hr before loading cells.

### RT-qPCR

Total RNA was isolated from cells using the PureLink RNA Mini Kit (Thermo Fisher; 12183018A) and was used for reverse transcription using a High-Capacity cDNA Reverse Transcription Kit (Thermo Fisher; 4368814). qPCR was performed using PowerUp SYBR Green Master Mix (Thermo Fisher; A25742) on a real-time PCR detection system (CFX96; Bio-Rad). Relative mRNA levels were calculated using the ΔCt method.

### RNA interference

Non-targeting (4390844), HMGCR (4392422; s141), Piezo1 (4392420; s18891), and MYPT1 (4390824; s9235) siRNAs were purchased from Thermo Fisher (Waltham, MA). All siRNA transfections were performed using RNAiMAX (5 µL; Thermo Fisher) and OptiMEM (400 µL; Thermo Fisher). 200,000 cells were trypsinized and seeded in 6-well plates in complete media. After cells adhered (∼1 hr), siRNAs in RNAiMAX/OptiMEM were added to cells in complete media (2 mL) at a final concentration of 50 nM. Cells were incubated with siRNAs for 3 days.

### Transmigration

12 µm (VWR; 10769-224), 8 µm (Corning; 3422), or 5 µm (Corning; 3421) transmigration filters were incubated with fibronectin (10 µg/mL) (Thermo Fisher; 33016015) for 12 hr at 4 °C. After drying for 30 min and washing with PBS, cells were seeded at a density of 20,000/cm^2^ in starve media with BSA (0.1 %) into the upper well. Media containing fetal bovine serum (FBS; 10%) (Sigma Aldrich; 12106C) was added to the lower well. Both the upper and lower wells were treated as indicated. After 12 hr, transmigrated cells were stained, imaged, and counted in Fiji (https://fiji.sc/).

### Proliferation

20,000 cells were seeded in 24-well tissue culture plates (Corning; 353047), treated as indicated, and cultured for 5 days. Each day, cells were lifted, resuspended in complete media, diluted 1:1 with Trypan Blue (Sigma; 8154), and loaded into cell counting slides (Bio-Rad; 1450011) for quantification using the Automated Cell Counter (Bio-Rad; 1450102).

### Intracellular cholesterol and LDL uptake

Following the instructions provided by the manufacturer, we used kits to measure intracellular cholesterol (Thermo Fisher; A12216) and LDL uptake (Thermo Fisher; I34360).

### Western blotting

Lysates were prepared on ice using a rapid lysate preparation kit (Cytoskeleton, Denver, CO; BLR01) in the presence of protease and phosphatase inhibitors (Cell Signaling Technology; 5872S). Samples were separated on gradient gels, transferred to nitrocellulose, and proteins were immobilized by air drying overnight before blocking (LI-COR, Lincoln, NE; 927-66003). Primary antibodies for pRLC (3671), RLC (8505), and GAPDH (5174) were purchased from Cell Signaling Technology (Danvers, MA) and diluted (1:500) in Tris-buffered saline containing 0.1% Tween 20 (TBS-T). Bands were resolved using infrared (IR) dye conjugated secondary antibodies on an Odyssey imager (LI-COR).

### 2D confinement

This protocol has been described in detail elsewhere (19). Briefly, PDMS (24236-10) was purchased from Electron Microscopy Sciences (Hatfield, PA). 2 mL was cured overnight at 37 °C in each well of a 6-well glass bottom plate (Cellvis; P06-1.5H-N). Using a biopsy punch (World Precision Instruments; 504535), an 8 mm hole was cut, and 3 mL of serum-free media containing 1% BSA was added to each well and incubated overnight at 37 °C. After removing the serum-free media containing 1% BSA, 300 μL of complete media containing trypsinized cells (250,000 to 1 million) and 2 μL of 3.11 μm beads (Bangs Laboratories, Fishers, IN; PS05002) were then pipetted into the round opening. The vacuum created by briefly lifting one side of the hole with a 1 mL pipette tip was used to move cells and beads underneath the PDMS. Finally, 3 mL of complete media was added to each well. Cells recovered for ∼60 min before imaging.

### Ca^2+^ measurements

For measurements of intracellular [Ca^2+^], cells were loaded with both a green fluorescent dye (Thermo Fisher; C7025) and a red fluorescent Ca^2+^ indicator, Calbryte (Fisher Scientific; NC2111763), for ratiometric fluorescence imaging.

### Microchannel preparation

PDMS (Electron Microscopy Sciences; 24236-10) was prepared using a 1:7 base and curing agent ratio. Uncured PDMS was poured over the wafer mold, placed in a vacuum chamber to remove bubbles, moved to a 37 °C incubator, and left to cure overnight. After curing, small PDMS slabs with microchannels were cut using a scalpel, whereas cell loading ports were cut using a 0.4 cm hole punch (Fisher Scientific; 12-460-409).

For making PDMS-coated cover glass (Fisher Scientific; 12-545-81), 30 µL of uncured PDMS was pipetted into the center of the cover glass, placed in a modified mini-centrifuge, and spun for 30 sec for even spreading. The PDMS-coated cover glass was cured for at least 1 hr on a 95 °C hot plate. To bond the slab and coated cover glass, PDMS surfaces were activated for ∼1 min by plasma treatment (Harrick Plasma; PDC-32G). The apparatus was incubated at 37 °C for at least 1 hr for complete bonding.

### Microchannel coating

Before microchannel coating, surfaces were first activated by plasma treatment. VCAM-1 (R&D Systems; 862-VC) or BSA (VWR; VWRV0332) was used for coating at 1 µg/mL and 1%, respectively, in PBS. Immediately after plasma treatment, VCAM-1 or BSA solution was pumped into microchannels using a modified motorized pipette. To remove any bubbles pumped into microchannels, the apparatus was left to coat in a vacuum chamber for at least 1 hr. Microchannels were then rinsed by repeatedly pumping in new PBS. Finally, microchannels were incubated overnight at 4 °C in complete media before use.

### Microchannel loading

Before cells were loaded into microchannels, complete media was aspirated, and microchannels were placed into an interchangeable cover glass dish (Bioptechs; 190310-35). Freshly trypsinized cells in 300 µL of complete media, stained with 1 µL of fluorescent membrane dye (Thermo Fisher; C10046), were pumped into microchannels using a modified motorized pipette. Once at least 20 cells were observed in microchannels by low magnification brightfield microscopy, microchannels were covered with 2 mL of complete media. Before imaging, a lid was placed on the apparatus to prevent evaporation.

### Live imaging

High-resolution live imaging was performed using a DeltaVision (Issaquah, WA) Elite imaging system mounted on an Olympus (Tokyo, Japan) IX71 stand with a motorized stage (XYZ), Ultimate Focus, solid state light source, fast filter wheel for DAPI, CFP, FITC, GFP, YFP, TRITC, mCherry, and Cy5 channels, critical illumination, Olympus UPlanSApo 20X/0.75 NA DIC (air), UPlanSApo 30X/1.05 NA DIC (silicone), PlanApo N 60X/1.42 NA DIC (oil), UPlanSApo 60X/1.30 NA DIC (silicone), and UPlanSApo 100X/1.40 NA DIC (oil) objectives, Photometrics (Tucson, AZ) CoolSNAP HQ2 camera, SoftWoRx (Preston, UK) software with constrained iterative deconvolution, cage incubator, and vibration isolation table.

### Genomics

Transcriptomic data for enzymes within the cholesterol biosynthetic pathway were obtained by The Cancer Genome Atlas (TCGA; https://www.cancer.gov/ccg/research/genome-sequencing/tcga) for skin cutaneous melanoma (SKCM). Data was sourced from the Genomic Data Commons (GDC; https://portal.gdc.cancer.gov/) and accessed through cBioPortal (https://www.cbioportal.org/). For survival curves, median mRNA levels (transcripts per million; TPM) for each enzyme were used as cut-off values.

### Cell classification in 2D confinement

Cells that displayed directionally persistent migration over at least 4 frames (32 min) were classified as leader mobile (LM). Any cell with a large bleb that remained stable for at least 4 frames (32 min) was considered to have a leader bleb.

### Cell classification in microchannels

Cells that moved at least ½ of their original length over a 5 hr timelapse movie were considered motile. Cells that displayed only blebs were classified as amoeboid, whereas cells that displayed only actin-based protrusions were classified as mesenchymal. Cells with blebs and actin-based protrusions were classified as hybrid.

### Morphometric analysis

The Fiji (https://fiji.sc/) plugin, Analyze_Blebs (https://github.com/karlvosatka/analyze_blebs), was used to measure the largest bleb area, aspect ratio, roundness, Feret’s diameter, solidity, and circularity of cells from timelapse movies (40).

### Cell migration

To perform cell speed and directionality ratio analyses, we used an Excel (Microsoft) plugin, DiPer, developed by Gorelik and colleagues, and the Fiji plugin (https://fiji.sc/), MTrackJ, created by Erik Meijering for manual tracking (41). Brightfield imaging confirmed that beads or debris were not obstructing the cell.

### FRET

Ratio images of FRET (CFP excitation/YFP emission) to CFP (CFP excitation/CFP emission) were generated and analyzed in Fiji (https://fiji.sc/).

### Statistics

Sample sizes were determined empirically and based on saturation. Statistical significance was determined using either a hypothesis test statistic or one-way ANOVA followed by appropriate multiple comparison tests in GraphPad (San Diego, CA) Prism. Outliers were identified using the ROUT method (Q = 1%). Normality was determined by a D’Agostino&Pearson test. For categorical data, χ2 tests were used to determine statistical significance. Significance levels: * - p ≤ 0.05, ** - p ≤ 0.01, *** - p ≤ 0.001, and **** - p ≤ 0.0001

## VIDEOS

**Video S1**. A vehicle treated melanoma cell confined to ∼3 µm by a PDMS ceiling coated with BSA (1%) adopt a leader mobile (LM) phenotype.

**Video S2**. A Fluvastatin treated melanoma cell confined to ∼3 µm by a PDMS ceiling coated with BSA (1%) adopt a no leader (NL) phenotype.

## SOURCE DATA

**Source data 1**. Categorical data plotted in Figures 1E, 1I, 1K, 5B, 5F, and S1A and associated χ^2^ tests.

## SUPPLEMENTAL INFORMATION

**Supplemental Figure 1.**
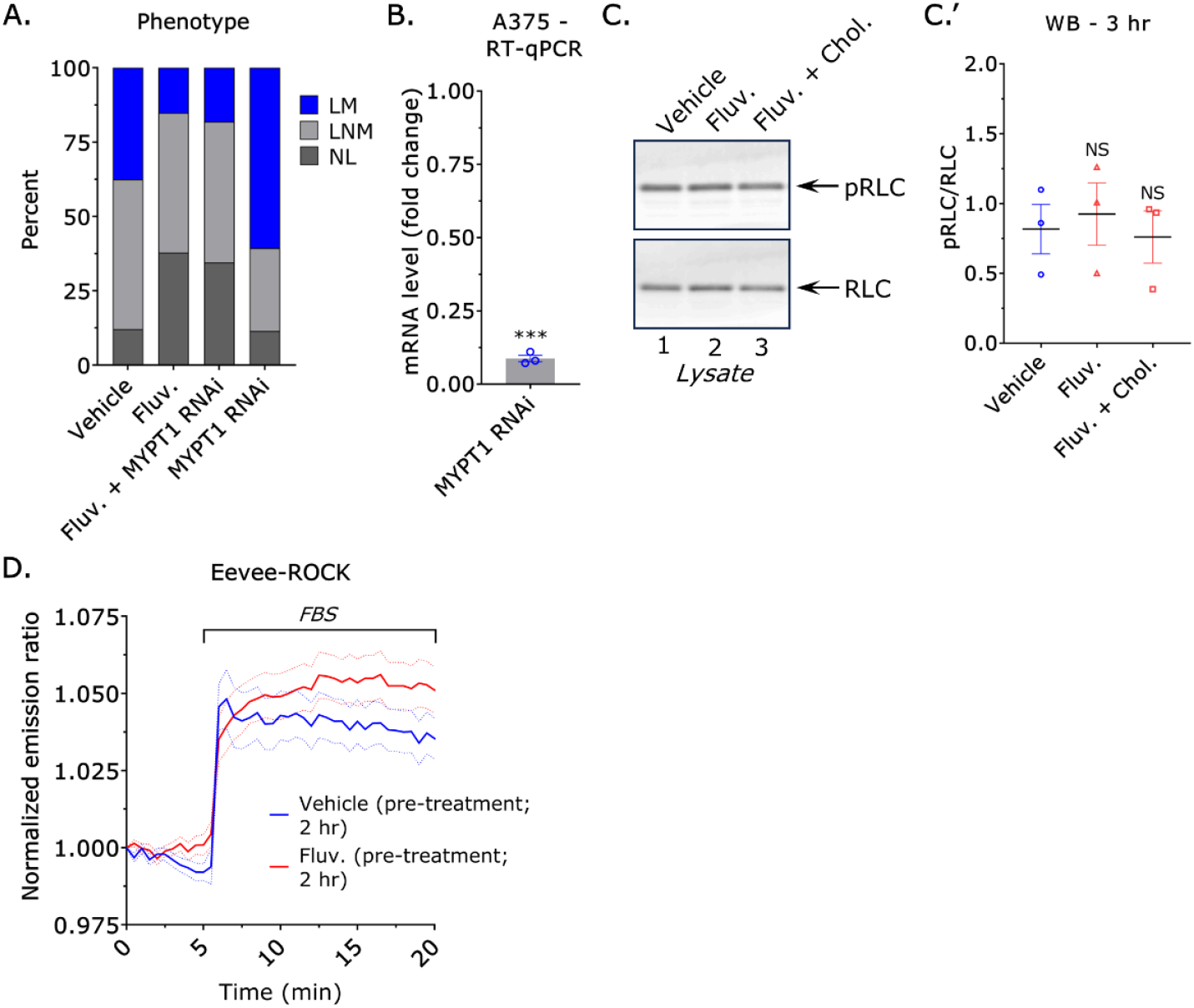
Cholesterol depletion does not impair actomyosin contractility. Percentage of leader mobile (LM), leader non-mobile (LNM), and no leader (NL) for vehicle (*n* = 334), Fluvastatin (10 µM; *n* = 340) (χ^2^ < 0.0001), Fluvastatin (10 µM) + MYPT1 RNAi (*n* = 530) (χ^2^ < 0.0001), and MYPT1 RNAi (*n* = 320) (χ^2^ < 0.0001) cells. **B**. Fold change in mRNA after MYPT1 RNAi, as measured by RT-qPCR (mean +/- SEM). **C**. Western blots against phosphorylated regulatory light chain (pRLC; S19) and regulatory light chain (RLC) from lysates of vehicle, Fluvastatin (10 µM; 3 hr), and Fluvastatin (10 µM; 3 hr) + cholesterol (1 mM; 3 hr) treated cells (mean +/- SEM). A Dunnett’s multiple comparison test was used to determine statistical significance. **D**. Normalized emission ratio for Eevee-ROCK from starved cells pre-treated (2 hr) with vehicle or Fluvastatin (10 µM). Fetal bovine serum (FBS; 10%) was added after 5 min (mean +/- SEM). Significance levels: * - p ≤ 0.05, ** - p ≤ 0.01, *** - p ≤ 0.001, and **** - p ≤ 0.0001. See also source data 1.

